# Theoretical value of radiation priming for mitigating toxicity in radiation therapy

**DOI:** 10.1101/2025.08.15.670485

**Authors:** Yair Y. Shaki, Yehoshua Socol

## Abstract

The relevance of ionizing radiation effects is high in the field of radiation oncology. Namely, radiation toxicity is a crucial limiting factor in dose administration. To address this concern, by adopting an empirical approach it is possible to uncover certain truths that might not be attainable through rigorous biophysical analysis, as the human body represents a complex system. That is, even possessing complete knowledge of interactions between its elementary units (cells) may prove insufficient in explaining the overall system’s reactions. An empirical mathematical model for adaptive biological response to time-dependent irradiation of arbitrary time-shapes was developed. The model was based on the damped-oscillator analogy from physics. The research reported here is a generalization of the damped-oscillator model by introducing a biologically relevant inverted Gompertzian-like shape, rather commonly employed in adaptive response modeling, with practical implications for biologists and oncologists. Based on the developed model, a practical approach is suggested to improve radiation therapy protocols by radiation training (priming) through administration of low, gradually increasing total-body irradiation doses. Following this priming, individuals should be capable of withstanding higher therapeutic doses, maybe up to five times the regular amount. Initial estimations propose a training time of approximately six weeks and a training dose of up to approximately 600 cGy. If the model is correct, effectiveness of radiation therapy can be significantly improved by radiation priming, since administration of considerably higher therapeutic doses would be enabled through priming procedures.

## 1. Background

### 1.1 Introduction

The impact of ionizing radiation is high in the field of radiation oncology. Radiation therapy (RT) is a prevalent treatment modality, employed in approximately 60% of cancer patients [1]. However, radiation toxicity poses a critical constraint on the therapeutic dosage, and despite substantial technological advancements, the adverse effects of RT remain considerable [2]. Of particular interest is the potential for incorporating relatively low radiation doses as an adjunctive therapy. This concept was initially explored by Sakamoto et al., who achieved successful lymphoma therapy [3], and more recently, by Kojima et al., who conducted case studies involving cancer and non-cancer diseases [4]. Both groups’ research demonstrates that low doses can be beneficial as complementary treatment augmenting the body’s response, despite the fact that relevant mechanisma are far from being fully understood.

Another crucial question pertains to the possibility of enhancing the patient’s radioresistance. Greater radioresistance in healthy tissues (and the organism in general) could prove pivotal in radiotherapy, as it may mitigate adverse effects when employing higher doses to target tumors. This prospect can be traced back to Yonezawa’s seminal work, which demonstrated an adaptive response, or hormesis, through experiments conducted on a mouse model [5]. The study involved exposing a subcohort of mice to low-dose irradiation (priming) before administering a high radiation dose. The important finding was that the survival rate of mice subjected to preliminary irradiation (priming) was significantly higher compared to those without priming. Smirnova and Yonezawa [6] subsequently proposed a mathematical model to elucidate the abovementioned experimental findings. However, the model was exceedingly intricate with 13 first-order differential equations and approximately 50 parameters. In light of this complexity, Fornalski et al. [7] devised a novel theoretical approach to comprehend and explain this phenomenon. Their study specifically examined separate detrimental effects.

The human body is a complex system: it exhibits intricate dynamics, where an exhaustive understanding of interactions between elementary units may fall short in explaining the system’s overall response. Therefore, adopting a purely empirical approach to describe and model these processes could potentially unveil truths not readily apparent when solely analyzing the biophysics of cells. Analogies with rather simple physical systems may be important.

### 1.2 Low-dose radiation response as a damped oscillator evolvement

Another mathematical model describing the biological response to time-dependent irradiation of arbitrary time-shape has been developed. The model [8], [9] draws an analogy with a damped oscillator. The concept of damped oscillator in physics refers to a oscillating system with oscillations decay because of damping. Some examples are a mass on a spring, or a swinging pendulum. This model quantifies the organism’s resistance to acute radiation effects through a single characteristic, which can be called “resistance to illness” or “organism strength,” which may relate to parameters such as hematopoietic proliferation rates and antibody presence. However, rather than relying on these associations, the problem has been treated phenomenologically. Namely, we aim to explain the following intuitive assumptions via a model, without insisting on building a complete mathematical theory.

First, we define two formal parameters u and τ. Their correspondence to physical parameters is discussed later in sec. 1.3.

- u – dimensionless excess of organism’s strength. The zero value u = 0 is defined as corresponding to the initial organism’s strength before irradiation, and u = x corresponds to x-fold excess of the strength, i.e., (x+1)-fold radiation resistance of the organism. For example, u = 3 corresponds to four-fold radiation resistance.
- τ – dimensionless time measured in units of the characteristic time T_0_.

The model assumes the following regarding the organism’s strength evolution over time:

- The time evolution of the organism’s strength is mathematically analogous to a mass on a spring with a damper (Fig. 1A). Following displacement from equilibrium, resulting from the acute irradiation, the organism undergoes one oscillation with so-called time constant (characteristic time) T_0_ as seen in Fig. 1B, and subsequently, the organism’s strength asymptotically approaches equilibrium. Such one-oscillation evolution is referred to as “critical damping.”
- Following acute irradiation (radiation pulse), the organism’s strength experiences an initial drop (degradation) proportional to the radiation dose.
- The organism’s response after acute irradiation is also proportional to the radiation dose. While both the response and strength degradation exhibit proportionality to the initial dose, the assumption of critical oscillation suggests that the strength eventually surpasses its pre-irradiation level.
- Upon subsequent radiation pulses of the same dose, both the strength and its time-derivative exhibit similar discontinuities as observed after the first pulse (Fig. 1C).

**Figure 1.**
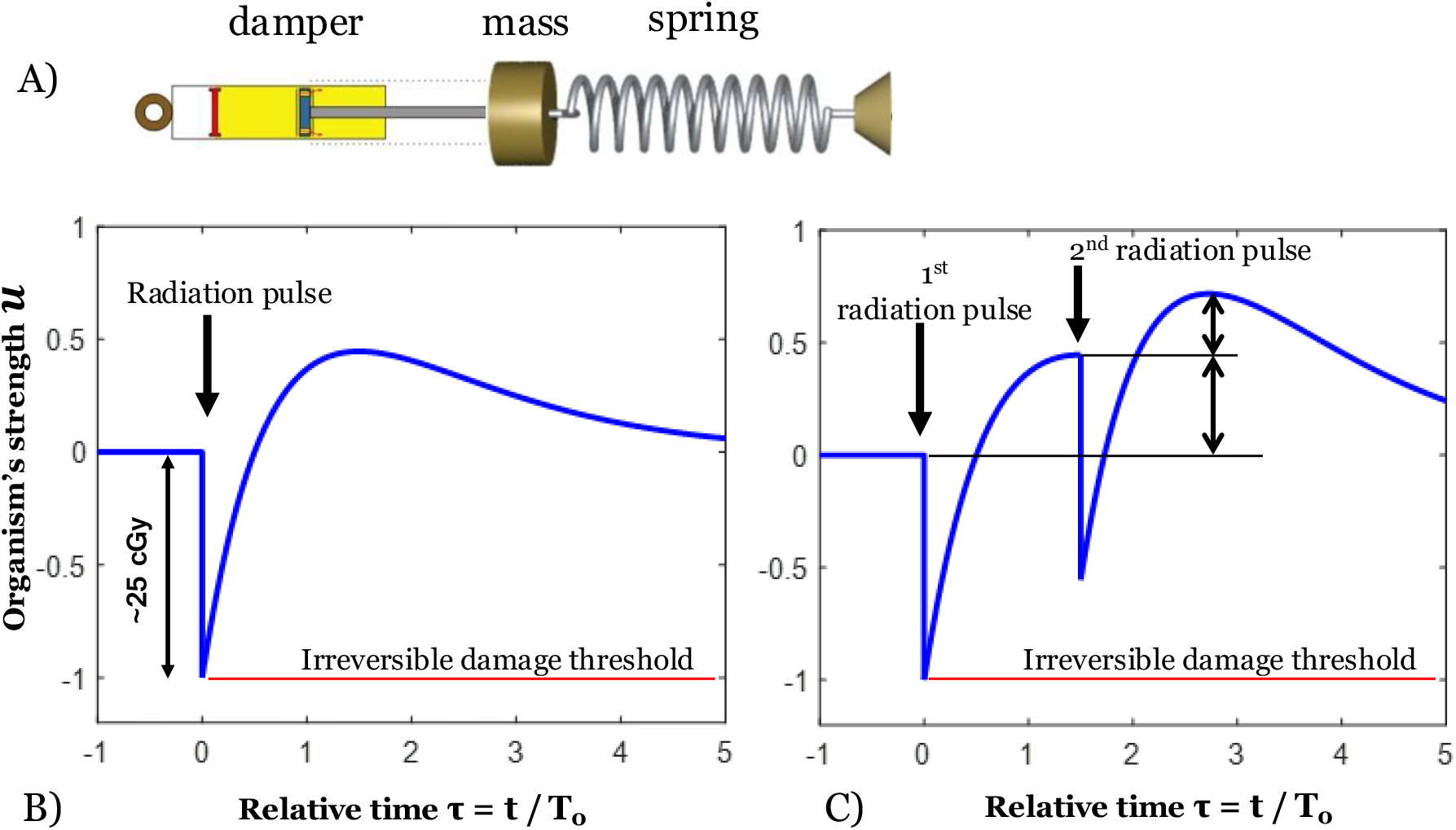
The damped oscillator mechanical analogy (A) and time evolution of the organism’s strength described by the model after single pulse (B) and double-pulse (C) irradiation. According to the proposed model, if we increase the irradiation dose is increased above about 25 cGy, the body will suffer irreversible damage. T_0_ is the time constant, or characteristic time of the system, hypothesized to be of the order of two weeks.

Under the premise of the aforementioned assumptions, the prospect of enhancing a patient’s radioresistance emerges. The enhancement is expected to be achieved through the implementation of “radiation training” involving a well-coordinated sequence of acute irradiations (fractions). Figure 1 exemplifies the impact of such fractionation, demonstrating that after the second pulse, the organism’s strength *u* attains a maximum of 0.8: this corresponds to a 80% increase in the dose that can be administered to the patient without causing irreversible damage (elaborated upon in the subsequent section).

### 1.3 Description of the parameters

If the radiation dose is too high, the body will suffer irreversible damage, resulting in a permanent decrease in its ability to fight infections and other problems. We think that this critical radiation level, where such damage occurs, is close to the point where acute sub-clinical effects begin to show, which is D(damage)=25 cGy – see, e.g., [10]. Interestingly, this value is also observed as the threshold for cancer development when studying atomic bomb survivors – see [11]). However, it is important to note that the value of 25 cGy is just a rough estimate and should be only used to indicate general trends based on the model. In the rest of the paper we assume that D(damage)=25 cGy corresponds to *u* = −1 (see Fig. 1) and actually describe the radiation dose in the dimensionless units of the dose divided by D(damage).

In terms of time, we assume that the body’s ability to adapt to radiation takes about 2 weeks. This estimation is based on practical experience, as acute side effects of radiation therapy usually subside a few weeks after treatment [2]. This value (T_0_ ∼ 2 weeks) aligns well with what we know about the effects of Acute Radiation Syndrome (ARS) from the atomic bombings in Japan. Deaths began after about a week from those bombing, and most ARS-related deaths peaked around 3 weeks and declined by 6 to 8 weeks (see Ref. [12] page 28). This information is particularly valuable since the atomic bomb victims did not receive special treatment, making their outcomes a reflection of natural processes, unlike, for example, the Chernobyl victims.

The model’s parameters are expected to differ from one individual to another, but likely within reasonable ranges of several tens of percent. For instance, the US Department of Defense has reported that a radiation dose resulting in a 30% probability of experiencing nausea/vomiting (the initial stage of ARS) is between 75-125 cGy [13]. We should stress that the above value is much higher than the values usually cited for hormesis – about 10 cGy [14]: mechanisms related to radiation responses are completely different for low doses and high ones and cannot be simply extrapolated. In practical applications of the model to real cases, the parameters described below are anticipated to be personalized and established at the outset of each specific treatment.

## 2. Dose dependence of the adaptive response

The dose dependence of the adaptive response is crucial. Using the analogy with the mass-spring-damper system, it can be said in simple terms that the dose dependence describes how the spring loses its elasticity as being stretched. We do not know this dose dependence, but we hypothesize that the functional law does not vary a lot from person to person. In the work [9] three hypotheses for functional dependency were suggested, while none of them could be associated with reasonable radiobiological mechanism. Here, we propose a general form of such dependency based on inverted Gompertzian-like shape which is widely used in adaptive response modeling – see, e.g., Ref. [7]. Several other threshold functions (binary and sigmoid) have been also tried with no significant difference in results. The cumulative function was selected since it was compatible with the model, while its derivative (Gompertz curve) was not.

Mathematically, Gompertz distribution has two parameters. We prefer to parametrize it somewhat differently from the canonical form, choosing one paprameter to be “effective dose” when the adaptive response decreases to 50% (the “spring’s” elasticity goes down to 50% of the initial value). We denote this parameter as ED_50_ (in loose analogy with LD_50_). The second parameter is the canonical Gompertz parameter η which has probably no biological meaning but descibes the shape width. Fig. 2 presents inverted Gompertzian-like shapes for 4 values of η. More mathematical details are given in the Appendix; for a thorough description of the model the reader is referred to the works [8-9].

**Figure 2.**
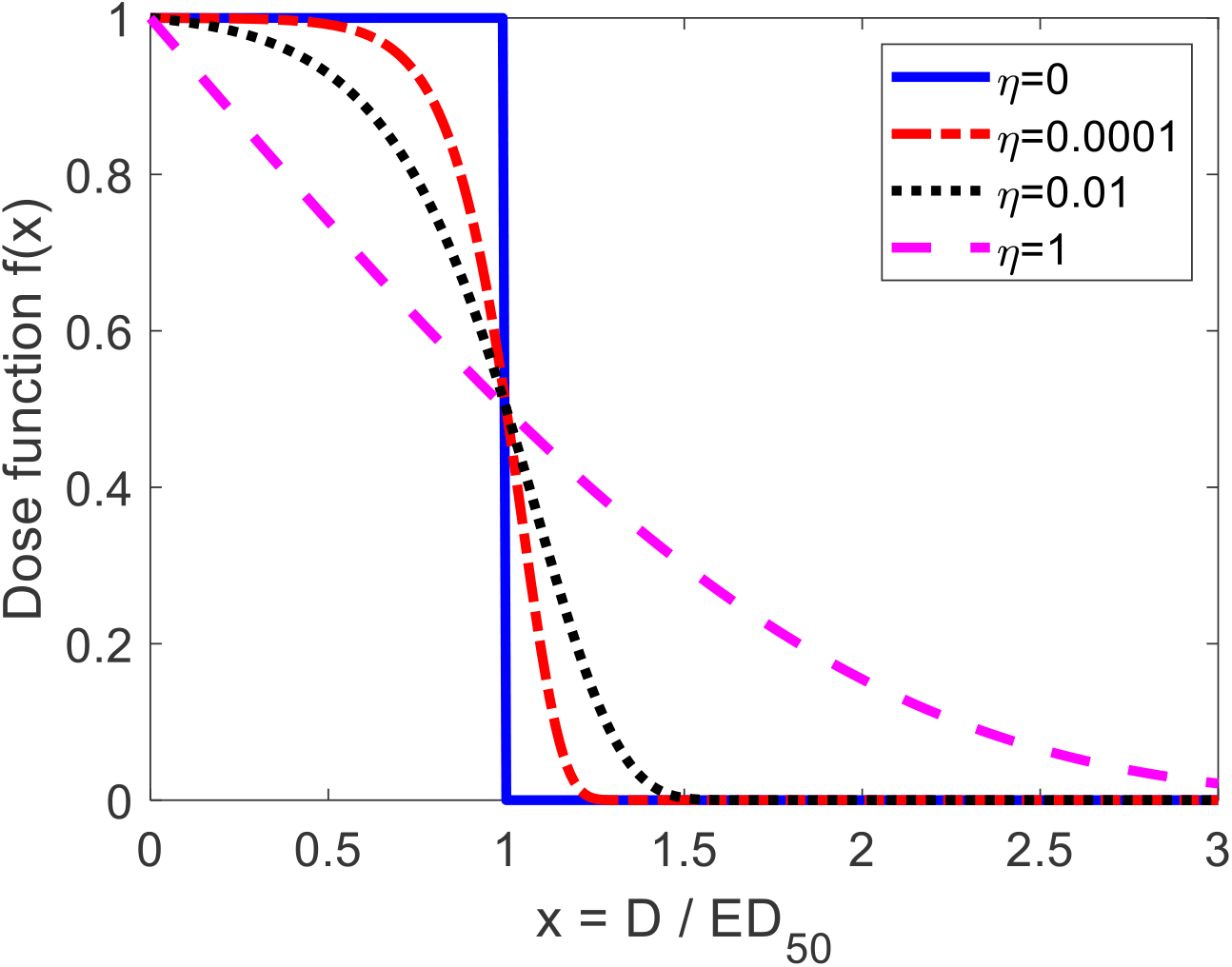
Four hypothetical dose functions of the organism’s adaptive response corresponding to four values of η of inverted Gompertzian-like shape. ED_50_ is “effective dose” when the adaptive response decreases to 50%. The parameter η has probably no biological meaning but descibes the shape width.

One should notice that the function Fig. 1 is “strength”, while the function in Fig. 2 is the dose dependence function. Dose dependence describes – mechanically speaking – how the spring loses its elasticity as being stretched.

## 4. Radiation training

As mentioned earlier in the introduction, the side effects of radiation therapy (RT) are significant. While dose fractionation is commonly employed to lessen these side effects, no kind of training has been reported for this purpose. With whole-body low-dose irradiation, the aim is to enhance the body’s resistance to radiation while not affecting the tumor’s resistance, as tumors receive much higher radiation doses during RT compared to healthy tissues.

In the context of our model, a crucial practical concern is achieving a high value of *u* = *u*^target^ without surpassing the “red line” of irreversible damage, as illustrated in Fig. 3 (*u* = −1). In practice, certain safety margins should be considered, and thus, the minimum allowable level of *u* should be set above the value of *u* = −1, corresponding to the previously mentioned D(damage).

**Fig. 3.**
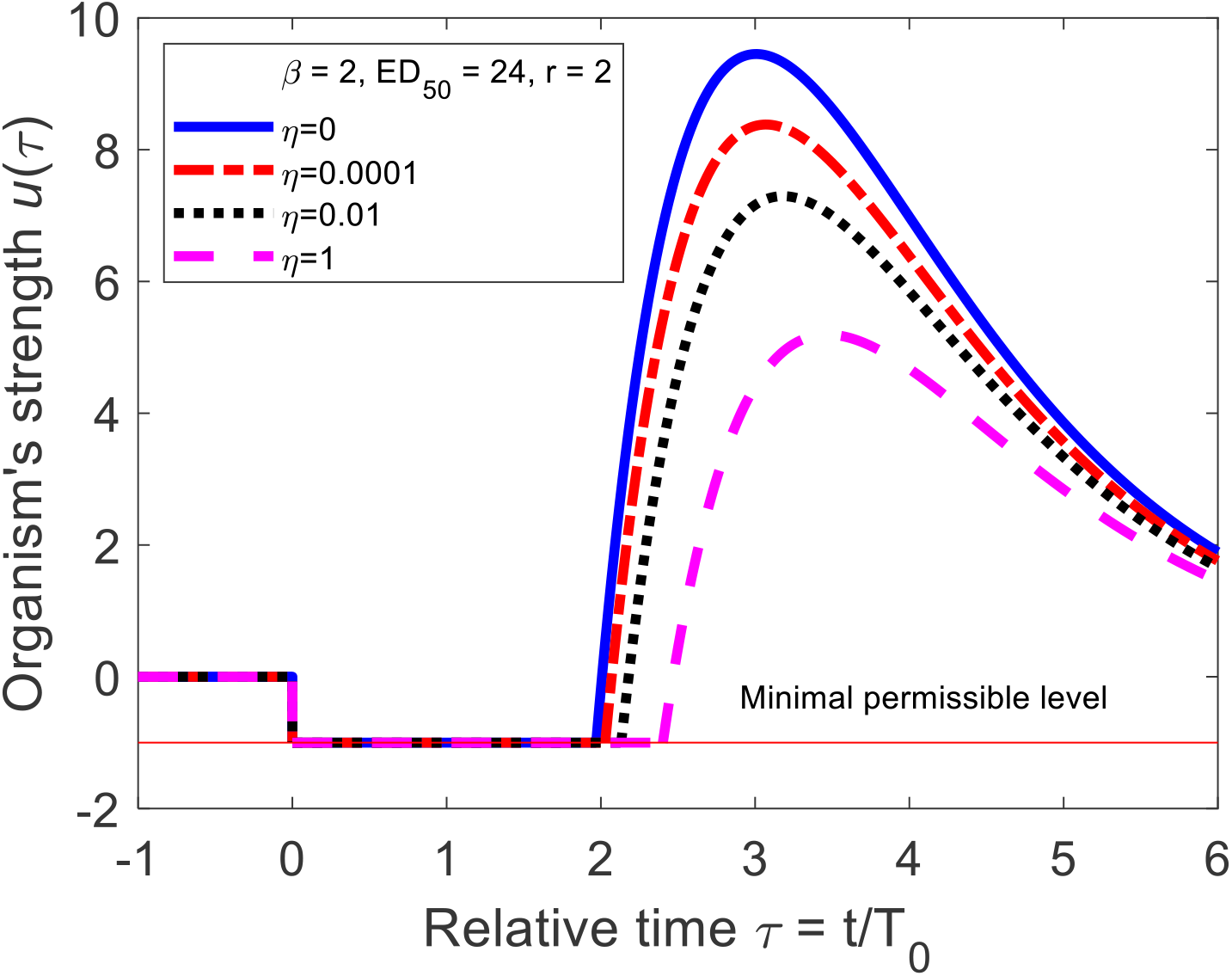
Values of the organism’s strength *u(*τ) as a result of the radiation training. The value *u(*τ) = 0 corresponds to the normal level. The value *u(*τ) = −1 corresponds to the minimal permissible level; normal person can sustand D(damage) (which we assume to be about 25 cGy) before the strength falls to *u(*τ) = −1. If *u(*τ) reaches the value of *u(*τ) = 1 then *u(*τ) + 1 = 2 and the patient is able to absorb radiation dose 2×D(damage) (twice of normal). The value *u(*τ) = 5 (η = 1) corresponds to 6-fold increase in radioresistance.

We propose the following heuristic approach as a solution. The suggested irradiation schedule involves three distinct stages:

1. Initial total-body irradiation with a relatively low dose, approximately 25 cGy.
2. Subsequent daily irradiation for τ_rad_ days, starting with even lower doses that increase exponentially over time.
3. A treatment pause, during which we anticipate the development of an adaptive response, making the patient’s organism (but hopefully not the tumor) more resistant to radiation. While cancer cells have usually completely different radiosensitivity (which is a basis for radiotherapy!), we anticipate that the tumor does not become more radioresistant because adaptive response is a systemic feature of the whole organism, not of particular tissue or tumor. Our anticipation is also based on Sakamoto’s results [3] showing better clinical outcome of combing low-dose and high-dose irradiation as compared to purely high-dose.

To some extent, the system behaves like a spring being gradually stretched until τ<τ_rad_, and then released when the irradiation is halted at τ=τ_rad_ (the analogy is only qualitative). The outcomes are presented in Fig. 3.

In Fig. 3 we assumed the dimensionless dose parameter *ED*_50_ = 24 which is hypothized to correspond (see 1.3) to about 600 cGy of absorbed dose. We hypothize that the adaptive dose response is actually described by the values of η about 0.0001–0.01 since such curves (Fig. 2) have similar shape to invereted dose response of radiation carcinogenesis observed in the survivors of the atomic bombing of Japan – see sigmoidal fit in Ref. [11], Fig. 4. But even for η = 1, it might be possible to achieve above fivefold increase in the patient’s radioresistance (Fig. 3).

The optimal duration of such training is expected to be around two units of the characteristic time T_0_ (as illustrated in Fig. 3). Assuming T_0_ = 14 days would require 4 weeks of total-body (TB) irradiation, and an additional 2 weeks to allow for the buildup of adaptive protection. (The question of whether delaying the RT treatment by 6 weeks is clinically acceptable goes beyond the scope of this discussion). The estimated TB dose is approximately *ED*_50_, i.e. about 600 cGy as dicussed above.

In this context, it is worth mentioning the scenario presented by Sakamoto [3]. It should be stressed that while Sakamoto studied long-term cancer effects, we are focusing on short-term tissue reactions. Although he employed dose fractionation (10 individual doses with a total dose of 150 cGy) for different purposes, those 10 individual doses administered a few times a week could be seen as somewhat similar to the proposed training. Nonetheless, daily irradiation starting with even lower but gradually increasing doses appears to be a more suitable approach.

## 4. Summary

We propose modeling the time evolution of the adaptive protective response to radiation as that of a damped oscillator, akin to a mechanical system of a mass attached to a spring with a damper. The model utilizes only a few parameters, which were roughly estimated based on current knowledge of acute radiation effects in both humans and animals. However, these parameters should be determined more precisely through future experiments, including animal models and clinical trials.

Utilizing the presented mathematical model, we have suggested a practical approach to enhance effectiveness of radiation therapy. By gradually increasing total-body priming doses, a person who undergoes this radiation treatment is expected to be capable of tolerating higher therapeutic doses, probably up to fivefold. Our very preliminary estimate indicates that the training period would last about six weeks, with a total training dose of approximately 600 cGy.

However, it should be considered that the adaptive response mechanisms are only partially understood [15] and despite the existing evidence [3-6] one cannot be sure that the adaptive response phenomenon will really appear. But even if the model is accurate for a (considerable) fraction of cancer patients, the suggested approach has the potential to significantly improve the effectiveness of radiation therapy (for these patients) by enabling the administration of considerably higher therapeutic doses through such training. Further research and experimentation are crucial to validate and refine this theoretically promising technique.

**Appendix**

The cumulative distribution function of the Gompertz distribution is decribed by the formula

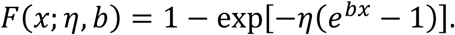

We use the inverted form

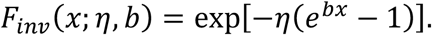

The function *F*_*inv*_*(x*; η, *b)* yields *f(x)* = 0.5 when

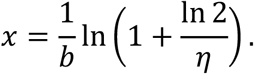

Therefore,

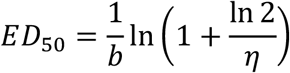

and

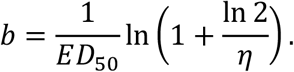

Correspondigly, taking *ED*_50_ and η as the two parameters instead of η and *b*, we can calculate *b* and *F*_*inv*_*(x*; η, *b)*.

